# Phosphoproteomics Maps Calcineurin-NFAT-DSCR1.4 Signaling as Druggable Axis in Gαq-R183Q–Driven Capillary Malformations

**DOI:** 10.1101/2025.06.07.658432

**Authors:** Tong Xu, Vera Janssen, Nathalie R. Reinhard, Paula Sobrevals-Alcaraz, Robert M. van Es, Annett de Haan, Julian de Swart, Martijn Wehrens, Albert Wolkerstorfer, Chantal M.A.M. van der Horst, Harmjan R. Vos, Stephan Huveneers

**Affiliations:** Amsterdam UMC, University of Amsterdam, Department of Medical Biochemistry, Amsterdam Cardiovascular Sciences, Meibergdreef 9, 1105 AZ Amsterdam, the Netherlands; Center for Molecular Medicine, Oncode Institute, University Medical Center Utrecht, Utrecht University, 3584 CG Utrecht, The Netherlands; Swammerdam Institute for Life Sciences, Section of Molecular Cytology, van Leeuwenhoek Centre for Advanced Microscopy, University of Amsterdam, Sciencepark 904, 1098 XH Amsterdam, the Netherlands; Amsterdam UMC, University of Amsterdam, Department of Dermatology, Amsterdam institute for Immunology and Infectious diseases, Meibergdreef 9, 1105 AZ Amsterdam, the Netherlands; Amsterdam UMC, University of Amsterdam, Department of Plastic, Reconstructive, and Hand Surgery, Amsterdam Cardiovascular Sciences, Amsterdam Public Health, Meibergdreef 9, 1105 AZ, Amsterdam, The Netherlands

**Keywords:** Sturge-Weber Syndrome, vascular malformation, endothelial cell, GNAQ p.R183Q, migration, angiogenesis, tacrolimus

## Abstract

Vascular malformations are congenital lesions caused by somatic and germline mutations that disrupt developmental signaling pathways. Capillary malformations (CMs) typically present as port-wine stains in the skin and can also affect ocular and cerebral tissues in Sturge Weber Syndrome (SWS), leading to aesthetic, ophthalmic, and neurological complications. CMs are caused by a somatic mutation in the *GNAQ* gene in endothelial cells, leading to a p.R183Q substitution in the Gαq protein. The underlying mechanisms of Gαq-R183Q-driven CMs formation remain unclear. To address this, we generated CRISPR/Cas9-engineered human dermal microvascular endothelial cells lacking endogenous Gαq, whilst expressing the Gαq-R183Q mutant instead. The Gαq-R183Q mutation strongly impaired endothelial cell migration and angiogenic sprouting capacity compared to wild-type controls. Next, using SILAC-based quantitative proteomics, we investigated the Gαq-R183Q-induced changes in the endothelial phosphoproteome. These analyses revealed prominent activation of the calcineurin-NFAT signaling pathway in Gαq-R183Q-expressing endothelial cells, leading to dephosphorylation of NFAT1 and NFAT2 and the selective expression of their transcriptional target DSCR1.4. Immunofluorescence of patient-derived skin biopsies confirmed deregulation of NFAT1/2 and the expression of DSRC1 in endothelial cells, validating their potential importance in CMs. We further demonstrate that pharmacological inhibition of calcineurin with tacrolimus (FK506) could partially restore NFAT signaling in Gαq-R183Q endothelial cells. Intriguingly, the genetic depletion of the NFAT target DSCR1 in Gαq-R183Q cells fully restored calcineurin/NFAT signaling to normal levels, enabling proper endothelial migration and sprouting. In summary, we uncovered a calcineurin-NFAT-DSCR1.4 signal transduction axis that is driven by Gαq-R183Q and established its importance for endothelial angiogenic properties. These findings highlight the calcineurin/NFAT signaling axis as a promising therapeutic target to restore endothelial function in CMs.

## Introduction

Vascular malformations are a group of congenital vascular and lymphatic lesions characterized by abnormally dilated and dysfunctional vessels. Based on the type of vessel involved, they are classified into venous, arteriovenous, lymphatic, and capillary malformations. Capillary malformations (CMs) occur in approximately 3 out of every 1,000 newborns and typically present as port-wine stains, which are marked by dilated and tortuous capillaries as well as post-capillary venules ^1^. A subset of patients with CM develops Sturge-Weber Syndrome (SWS), a neurocutaneous disorder in which a facial port-wine stain is accompanied by ocular and leptomeningeal capillary malformations^2^. Patients with SWS exhibit vascular hypertrophy and are affected by facial disfigurement, as well as neurological and ophthalmologic complications such as seizures, epilepsy, migraines, intellectual disability, and glaucoma ^1,3,4^. Current treatment options for SWS, including laser therapy, eye drops, and surgery to relieve intraocular pressure, are palliative rather than curative^1^. There is a clear need for better treatment strategies, and targeted molecular therapies offer a promising avenue by addressing the root causes of the vascular malformations^5^. Developing such treatments requires a thorough understanding of the molecular mechanisms behind CMs.

Both SWS and isolated port-wine stains are linked to a somatic mosaic mutation in the guanosine nucleotide-binding protein Q gene (*GNAQ*)^6–10^. This mutation results in an arginine to glutamine amino acid substitution at position 183 (p.R183Q) in the encoded Gαq protein (hereafter referred to as Gαq-R183Q)^7^. Sequencing studies have demonstrated that the Gαq-R183Q mutation is present in endothelial cells from multiple affected tissues, including: cutaneous endothelial cells of port-wine stains, brain endothelial cells in leptomeningeal malformations, and choroidal and scleral vessels in ocular lesions^8,9,11–15^. The Gαq protein and its close paralog Gα11 are α subunits of the heterotrimeric G protein complex, which serves as the primary signaling module downstream of G protein-coupled receptors (GPCRs). GPCRs are seven-transmembrane domain proteins that, upon binding to their cognate extracellular ligands, catalyze the activation of associated G proteins through GDP-GTP exchange on the Gα subunit.^16^. Upon activation, the GTP-bound Gαq generally activates phospholipase C-beta (PLCβ), which hydrolyzes the membrane phospholipid phosphatidylinositol 4,5-bisphosphate (PIP_2_) into two major second messengers: diacylglycerol (DAG) and inositol 1,4,5-trisphosphate (IP_3_). These second messengers then initiate multiple downstream signaling cascades, with DAG activating various protein kinase C (PKC) isoforms and IP_3_ triggering calcium release from intracellular stores^17,18^. The integrated activity of these pathways controls diverse cellular processes including growth, survival, proliferation, and migration ^19,20^. This signaling mechanism represents a widely conserved pathway across various cell types.

The specific downstream events of the Gαq-R183Q mutation in endothelial cells which contribute to CM formation are still unclear. Recent studies have demonstrated that Gαq-R183Q promotes constitutive PLCβ3 phosphorylation and subsequent activation of PKC and calcineurin signaling in endothelial cells, resulting in transcriptional changes^21^. Also the activation of Notch and MAPK/ERK pathways have been reported in Gαq-R183Q expressing endothelial cells^20,22^, however their contribution to disease development remains incompletely understood. In fact, a systematic analysis of protein signaling differences between healthy and Gαq-R183Q expressing endothelial cells and their functional contribution to CMs is currently lacking.

To develop targeted therapies for patients with port-wine stains and SWS, it is key to identify which molecules are specifically activated downstream of Gαq-R183Q signaling and to select those that constitute druggable targets. In this study, we sought to characterize altered signaling pathways downstream of endothelial Gαq-R183Q. This requires an unbiased characterization of the proteome and phosphoproteome affected by endothelial Gαq-R183Q, ideally from patient tissues. However, the utility of primary endothelial cells from CM skin biopsies carrying somatic Gαq-R183Q mutations is limited by their non-immortalized nature and finite lifespan in culture^9^. To overcome these limitations for proteomic scale studies, we established CRISPR/Cas9-mediated Gαq-knockout human dermal microvascular endothelial cell lines. These Gαq-knockout cell lines were subsequently rescued by expression of either wild-type Gαq (Gαq-WT) or mutant Gαq-R183Q. By using stable isotope labeling with amino acids in cell culture (SILAC)-based quantitative mass spectrometry, we performed comparative analyses of the phosphoproteome in Gαq-WT versus Gαq-R183Q-expressing endothelial cells. This unbiased proteomic approach identified differential protein abundance and phosphorylation levels associated with the Gαq-R183Q mutation. These analyses highlighted the importance the calcineurin-NFAT1/2-DSCR1.4 signaling axis in Gαq-R183Q-driven vascular malformations. By performing functional perturbation studies we could show that inhibition of calcineurin or perturbation of DSCR1 expression restores key endothelial cell functions in a Gαq-R183Q mutated background. These findings advance our understanding of the cellular and molecular mechanisms underlying CM pathogenesis and will facilitate the development of targeted therapies in CM and SWS patients.

## Results

### Characterization of Gαq knock-out HDMECs rescued with Gαq and Gαq-R183Q

To investigate Gαq-R183Q specific signaling in endothelial cells, we first generated CRISPR/Cas9-mediated Gαq knock-outs from immortalized human dermal microvascular endothelial cells (HDMECs) using guides RNAs targeting *GNAQ*. Next, Gαq KO HDMECs were rescued with lentiviral expression of intramolecularly mTurquoise2-tagged Gαq-wild-type or Gαq-R183Q (Fig. 1A). Western blot analysis confirmed that both rescued cell lines (hereafter referred to as Gαq-WT and Gαq-R183Q HDMECs) expressed similar Gαq levels, which was approximately two-fold higher than the endogenous protein in the parental HDMECs (Fig. 1A, B). Immunofluorescence analysis of VE-cadherin and F-actin revealed that Gαq-R183Q expression led to cell elongation, but did not affect cell-cell junctions or cytoskeletal organization when compared to Gαq-WT HDMECs (Fig. 1C). Quantitative morphometric analysis confirmed that Gαq-R183Q HDMECs exhibited larger cell sizes relative to Gαq-WT controls (Fig. 1D).

**Figure 1.**
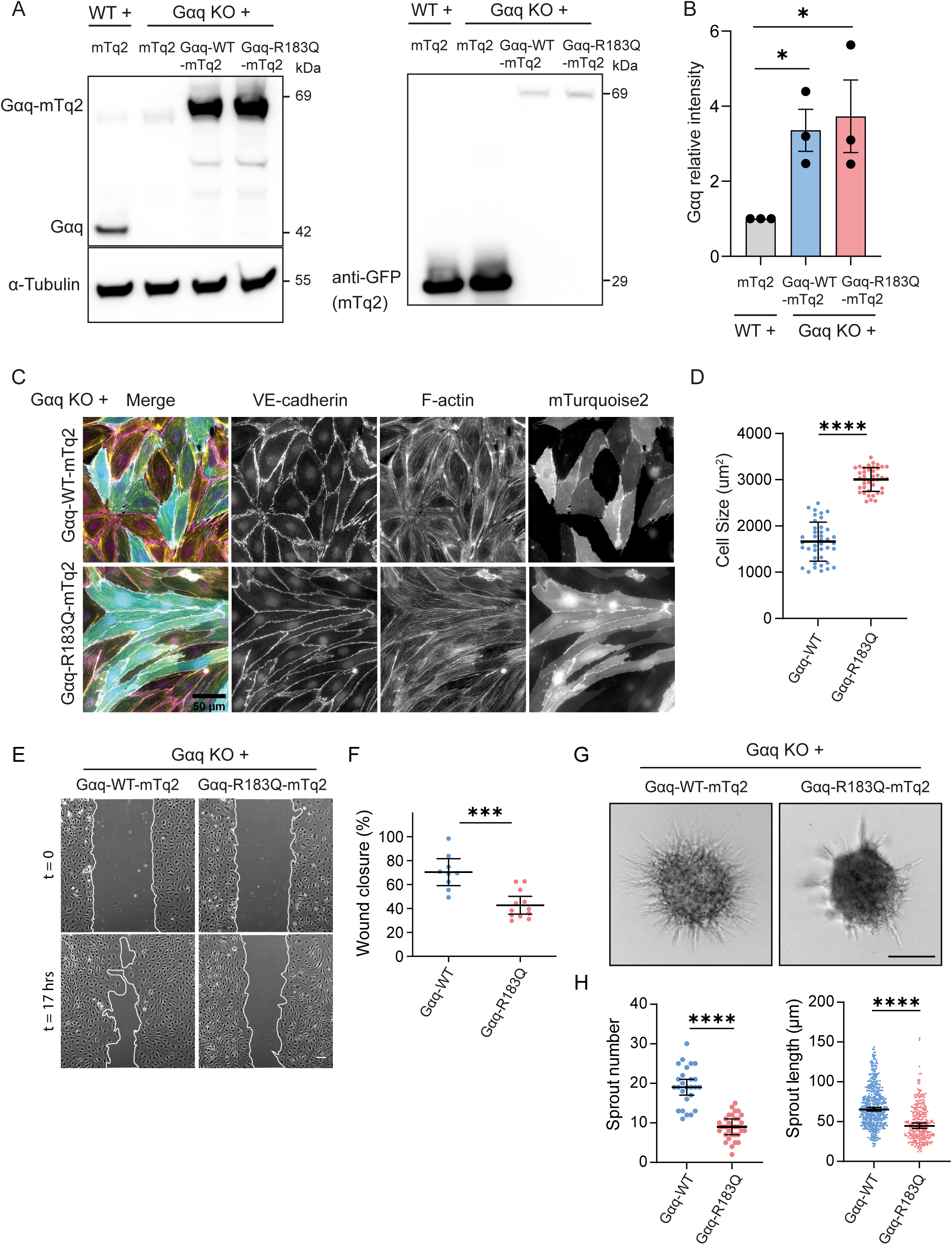
Impact of Gαq-R183Q mutation on endothelial cell morphology and function. (A) Western blot analysis of Gαq-WT-mTurquoise2 and Gαq-R183Q-mTurquoise2 re-expression in Gαq knockout (KO) human dermal microvascular endothelial cells (HDMECs). Left panel: Blots probed for Gαq and α-Tubulin (loading control). Right panel: Blots probed for GFP to detect the mTurquoise2 tag. (B) The bar plot shows quantification of Gαq band intensities. Data analyzed by one-way ANOVA with Tukey’s post hoc test; mean ± SD (n=3 independent experiments). (C) Widefield microscopy images of Gαq KO HDMECs rescued with Gαq-WT-mTurquoise2 or Gαq-R183Q-mTurquoise2. Cells were immunostained for VE-cadherin (magenta) and F-actin (yellow). Scale bar: 50 μm. (D) Cell area quantification in Gαq-WT-mTurquoise2 and Gαq-R183Q-mTurquoise2 HDMECs. Data analysed by one-way ANOVA with Tukey’s post hoc test; mean ± SD (n=5 independent experiments). Cell size for Gαq-WT mean =1661.078 μm^2^, SD =416.71 μm^2^, n = 40 cells, Gαq-R183Q mean =3004.675 μm^2^, SD =254.55 μm^2^, n = 40 cells, p = 0.0001. (E) Phase-contrast time-lapse images of scratch-wound assays in Gαq KO HDMECs expressing Gαq-WT-mTurquoise2 or Gαq-R183Q-mTurquoise2 at 17 hours post-scratch. White lines mark the unclosed wound area. Scale bar: 20 μm. (F) Dot plots show the percentage of remaining wound width. Data analysed by one-way ANOVA with Tukey’s post hoc test; mean ± SD (n=4 independent experiments; N = 9 Gαq-WT and 12 Gαq-R183Q samples). (G) Representative phase-contrast images of sprouting spheroids from Gαq-WT-mTurquoise2 and Gαq-R183Q-mTurquoise2 HDMECs. Scale bar: 50 μm. (H) Quantification of cumulative sprout number and length at 16 hours. Data analysed by one-way ANOVA with Tukey’s post hoc test; mean ± SD (n=4 independent experiments; N = 23 Gαq-WT and 27 Gαq-R183Q samples).

### Gαq-R183Q impairs endothelial cell migration and angiogenic sprouting

Since dysregulated endothelial cell migration and angiogenic sprouting are drivers of vascular malformations^23,24^, we investigated how Gαq-R183Q may affect these endothelial functions. We performed scratch wound assays of monolayers formed by Gαq-WT and Gαq-R183Q HDMECs. Gαq-R183Q expression significantly impaired endothelial cell migration toward the scratch resulting in delayed wound closure (Fig. 1E, F). To investigate the role of Gαq-R183Q in angiogenic sprouting, we conducted VEGF-induced spheroid-based sprouting assays. Sprout formation and elongation from spheroids was strongly decreased by Gαq-R183Q (Fig. 1G, H), confirming that proper Gαq signaling in ECs controls angiogenic behavior.

### Phosphoproteomics reveals differential activation of signaling pathways in Gαq-R183Q ECs

To investigate how the p.R183Q mutation in Gαq changes signal transduction, we performed SILAC-based quantitative phosphoproteomic profiling. Gαq-WT and Gαq-R183Q ECs were labeled with light or heavy amino acids (Lys-8, Arg-10), mixed post-lysis, and subjected to phosphopeptide enrichment. To address potential labeling biases, label-swap experiments, comprising forward and reverse replicates, were conducted (Fig. 2A). Phosphoproteomics on corresponding lysates identified differentially phosphorylated proteins (Table S1) and differentially abundant proteins (Table S2) in Gαq-R183Q ECs relative to Gαq-WT. Pathway mapping of all differentially phosphorylated sites using Ingenuity Pathway Analysis (IPA) revealed prominent upregulation of the phosphatase and tensin homolog (PTEN) pathway and downregulation of the integrin-linked kinase (ILK) pathway in Gαq-R183Q ECs (Fig. 2B). PTEN is a PI(3,4,5)P_3_ lipid phosphatase that antagonizes PI_3_K activity and suppresses Akt and mTOR signaling. Corresponding to the suppression phenotype, IPA regulator analysis showed reduced phosphorylation levels of PTEN effectors including AKT1, mTOR, and TSC1/3 in Gαq-R183Q ECs (Table S3). Somatic pathogenic mutations in *PTEN* were recently shown to disrupt PI3K/Akt/mTOR signaling in endothelial cells and drive overgrowth in vascular malformations^25,26^, highlighting the value of the Gαq-R183Q phosphoproteomic dataset for CM-relevant signaling.

**Figure 2.**
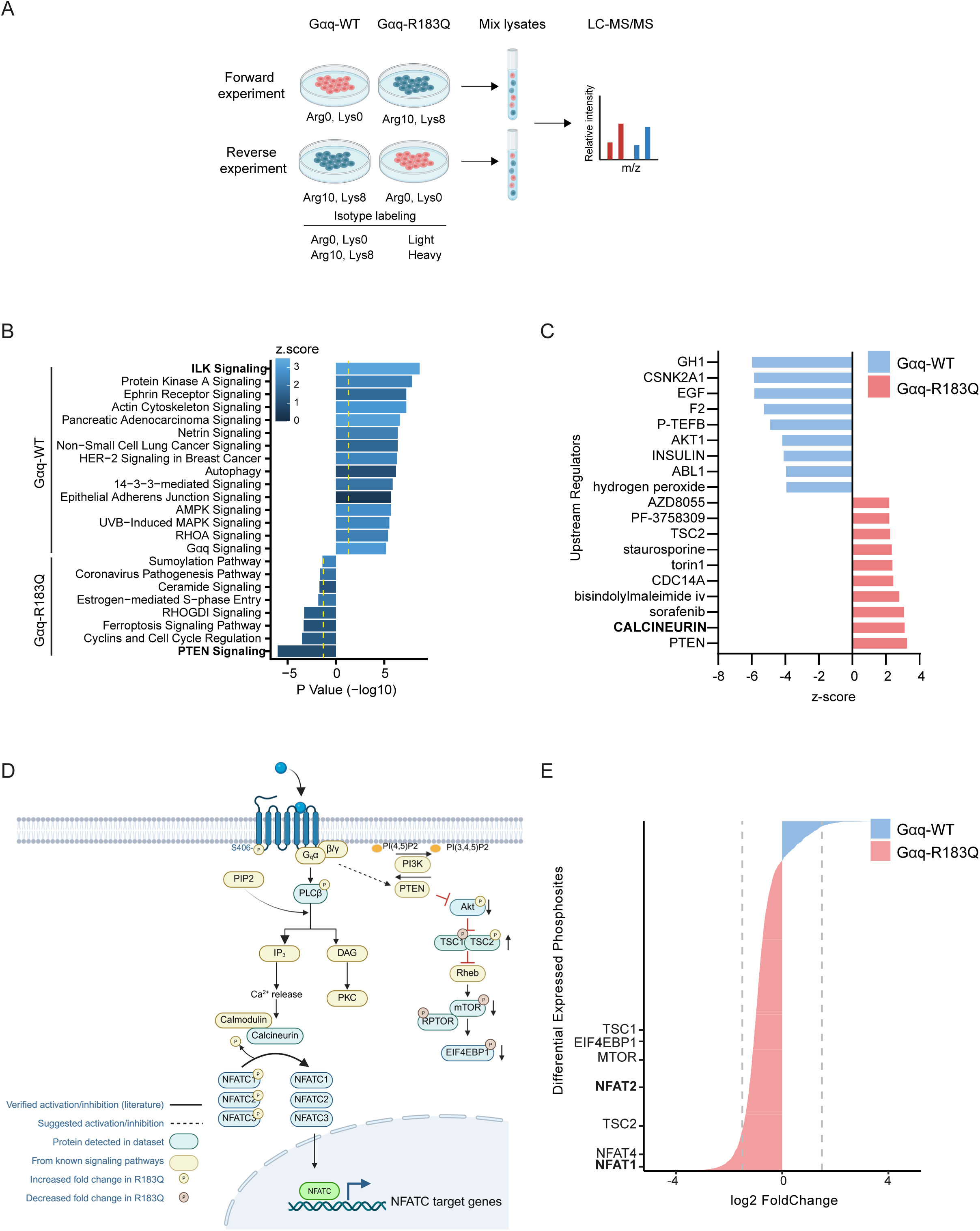
Phosphoproteomic profiling reveals differential signalling in Gαq-R183Q endothelial cells. (A) Schematic of the SILAC phosphoproteomics workflow. Stable isotope labelling of amino acids in cell culture (SILAC) was performed to compare Gαq-WT and Gαq-R183Q cells. Mixed cell lysates were processed for proteomic and phosphoproteomic analysis. Schematic created with Biorender.com. (B) Ingenuity Pathway Analysis to identify enriched signalling pathways for Gαq-WT and Gαq-R183Q. Top 15 significantly upregulated pathways for Gαq-WT and all Gαq-R183Q pathways are graphed according to their corresponding p-value and coloured according to the z. scores. Yellow dotted lines indicated thresholds with p < 0.041. (C) Ingenuity Pathway Analysis (IPA) of upstream regulators significantly enriched in Gαq-WT and Gαq-R183Q cells. All upstream regulators (for Gαq-WT) and top 10 significant upstream regulators (for Gαq-R183Q) are plotted by their activation Z-scores and -log(p-value) thresholds (p < 0.041). (D) Proposed Gαq-R183Q signalling pathway from integrating literature and SILAC phosphoproteomics data. Schematic created with Biorender.com (E) Fold change analysis of phosphosites of calcineurin targets between Gαq-WT and Gαq-R183Q. The dotted line represents the 1.5 absolute fold change cut-off.

To search for key kinases and/or phosphatases that could be responsible for the observed changes in phosphorylation, IPA upstream analysis was performed, identifying PTEN and calcineurin as the top hits in Gαq-R183Q ECs (Fig. 2C). Calcineurin is a known downstream effector of activated Gαq signalling. Upon activation, Gαq promotes PLCβ-mediated hydrolysis of PIP_2_, resulting in the production of IP_3_ and DAG. These second messengers initiate distinct signaling cascades: DAG activates PKC, while IP_3_ binds to its receptors on the ER membrane, triggering Ca²⁺ release into the cytosol (Fig. 2D). The resulting increase in intracellular Ca²⁺ facilitates the binding of calmodulin to calcineurin, thereby activating calcineurin-dependent signaling ^18^ ^27^ ^28^. Calcineurin, a serine/threonine phosphatase, dephosphorylates NFATC proteins^29,30^. Indeed, reduced phosphorylation levels of NFATC1/2/4 as well as other calcineurin targets were detected in the proteomics data (Fig. 2D, E). Taken together, the SILAC-based phosphoproteomic profiling of Gαq-R183Q endothelial cells revealed prominent PTEN pathway upregulation, ILK downregulation, and PLCβ-IP_3_-calcineurin activation, implicating altered signaling in *GNAQ*-related CMs.

### Disrupted NFAT1/2 phosphorylation levels, nuclear translocation and transcriptional activity in Gαq-R183Q ECs

The Gαq-R183Q-PLCβ-IP_3_ pathway has previously been associated with CM^31^, however the importance of its downstream signaling for endothelial function remains incompletely understood. Since our experiments indicated an important role for calcineurin-NFAT activation in Gαq-R183Q-ECs, we next investigated its regulation at molecular level. Generally, NFAT1 and NFAT2 dephosphorylation by calcineurin induces their nuclear translocation regulating gene transcription^32^ ^33^ ^34^. The SILAC analysis showed no difference in the abundance of total NFAT1 or NFAT2 expression in Gαq-WT and Gαq-R183Q ECs (Table S2), which we confirmed by Western blot analysis (Fig. 3A, B). The phosphoproteomics showed reduced phosphorylation levels at 4 distinct sites on the NFAT1 protein and 18 sites on NFAT2 in Gαq-R183Q ECs (Table S1). To validate these results, we performed Western blot analysis of phosphorylated Ser54 and Ser326 of NFAT1 and Ser294 of NFAT2. Indeed, phosphorylation levels of NFAT1-S54, NFAT1-S326, and NFAT2-S294 were significantly reduced in Gαq-R183Q cells (Fig. 3A, B). Intriguingly, immunofluorescence imaging experiments demonstrated that both NFAT1 and NFAT2 proteins remain in the cytoplasm of Gαq-R183Q ECs and fail to translocate to the nucleus like in Gαq-WT ECs (Fig. 3C, D). Automated quantitative analysis confirmed a strong decrease in nuclear-to-cytoplasmic ratio for both NFAT1 and NFAT2 in Gαq-R183Q cells (Fig. 3C, D). These results show that NFAT1 and NFAT2 proteins are constitutively dephosphorylated in Gαq-R183Q cells, however the majority of NFAT proteins remain in the cytoplasm, suggesting disrupted NFAT signaling. Next, we investigated protein expression of the NFAT target Down Syndrome Critical Region Protein 1 (DSCR1), which has been shown to be transcriptionally upregulated through overexpression of Gαq-R183Q in endothelial cells^31^. We detected an equal abundance of the canonical DCSR1.1 isoform, whereas levels of the DSCR1.4 isoform (also known as RCAN1.4), which acts as an inhibitor for NFAT through negative feedback signaling^35,36^, were strongly upregulated in Gαq-R183Q ECs (Fig. 3E, F). Together, these findings reveal that Gαq-R183Q promotes calcineurin-mediated NFAT dephosphorylation, followed by a negative feedback loop that limits nuclear NFAT translocation.

**Figure 3.**
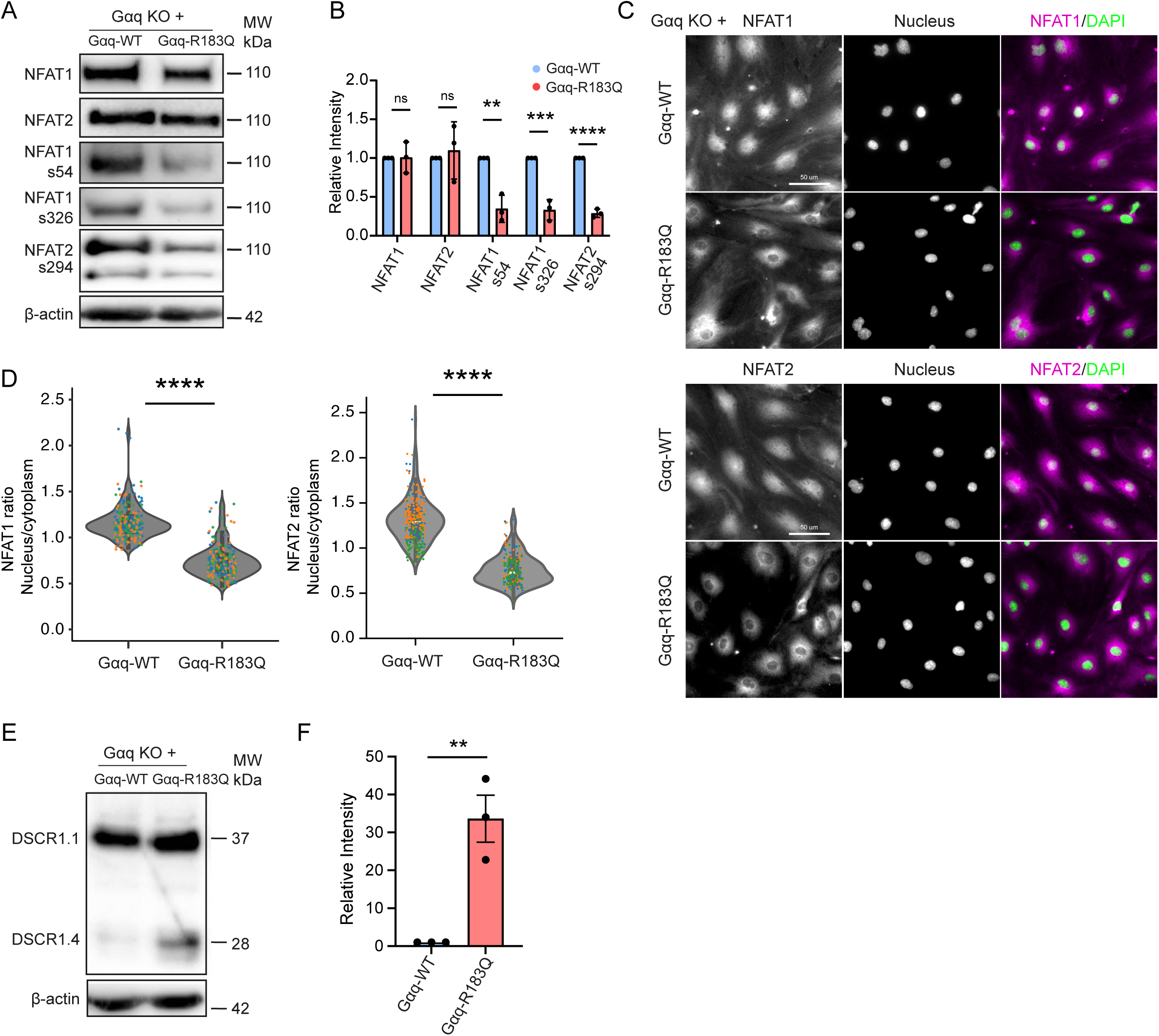
Disrupted NFAT1/2 phosphorylation levels, nuclear translocation and transcriptional activity in Gαq-R183Q ECs. (A) Western blot analysis of NFAT1, NFAT2, NFAT2-S294, NFAT1-S54, NFAT1-S326, and β-actin (top to bottom) in Gαq-WT and Gαq-R183Q cells. β-actin served as the loading control. (B) The bar plot shows quantification of band intensities. Data analysed by one-way ANOVA with Tukey’s post hoc test; mean ± SD (n=3 independent experiments). (C) Immunofluorescence images of Gαq-WT and Gαq-R183Q cells stained for NFAT1 (magenta, top), NFAT2 (magenta, bottom), and DAPI (blue; nuclei). Scale bar: 50 μm. (D) Quantification analysis of the nuclear-to-cytoplasmic ratio of NFAT1/2 fluorescence intensity using automated image processing and segmentation. Data analyzed by one-way ANOVA with Tukey’s post hoc test; mean ± SD (n=5 independent experiments; N= 225 Gαq-WT and 258 Gαq-R183Q cells for NFAT1; N= 208 Gαq-WT and 344 Gαq-R183Q cells for NFAT2). (E) Western blot analysis of DSCR1.1 and DSCR1.4 expression in Gαq-WT and Gαq-R183Q cells. β-actin was used as the loading control. (F) Data analysed by one-way ANOVA with Tukey’s post hoc test; mean ± SD (n=3 independent experiments).

### Characterization of NFAT1/2 and DSCR1 in skin capillary malformations harboring the *GNAQ* p.R183Q mutation

To validate whether disrupted NFAT signaling occurs within the endothelium of patients with CMs, we next performed immunofluorescence imaging of patient-derived skin biopsies that harbor the *GNAQ* p.R183Q mutation^9^. Immunostainings show DSCR1 protein expression in endothelial cells (marked by VE-cadherin) from CMs patients, as well as in epidermal epithelial cells (Fig. S1A). Immunostainings for NFAT1/2 indicated an enriched endothelial distribution and showed that both NFAT1 and NFAT2 proteins were predominantly localized in the endothelial cytoplasm, rather than in the nucleus (Fig. 4A, B). Of note, immunostainings using phosphorylated NFAT1-S326 and NFAT2-S294 did not result in positive signal in endothelial cells from CM patients (Fig. S1B). These results are consistent with the previous in vitro findings and suggest that NFAT signaling is disrupted in CMs.

**Figure 4.**
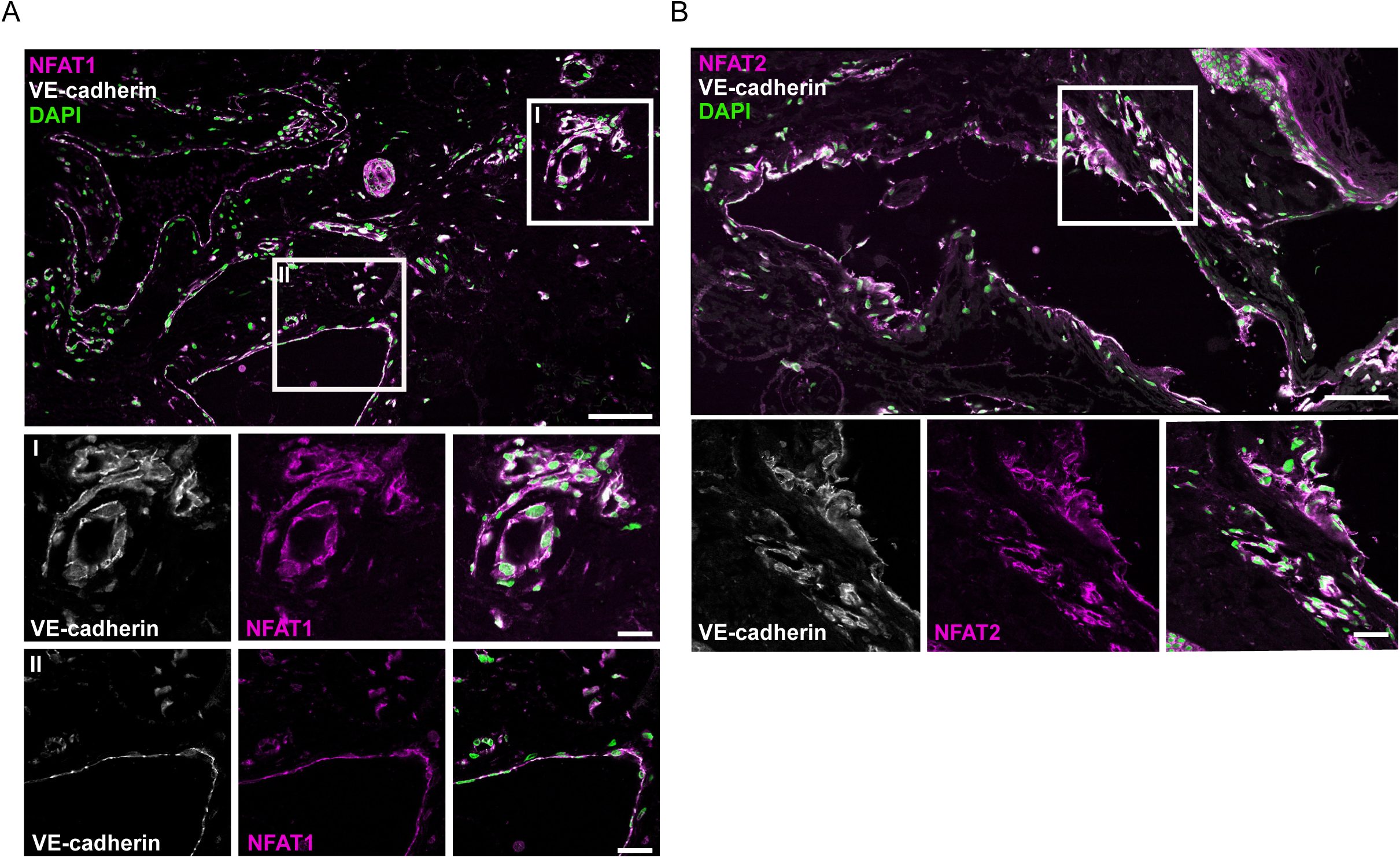
Characterization of NFAT1/2 and DSCR1 in skin capillary malformations harboring mutated a *GNAQ* p.R183Q mutation. (A) Representative confocal images showing NFAT1 (magenta) staining in skin biopsies harboring GNAQ p.R183Q mutation. Endothelial cells were marked with VE-cadherin (white), and nuclei were counterstained with DAPI (green). Scale bar = 100 µm. (B) Representative images showing NFAT2 staining in skin biopsies. Endothelial cells were marked with VE-cadherin, and nuclei were stained with DAPI. Scale bar = 100 µm.

### Inhibition of calcineurin activity improves functions of Gαq-R183Q ECs

Tacrolimus (FK506) is an immunosuppressant drug that mainly works by binding to FKBP12 which inhibits calcineurin activity^37^. To investigate the effect of FK506 on Gαq-R183Q-induced cellular events, we treated Gαq-R183Q ECs with 20 ng/ml FK506 for 24 hours. These treatments resulted in a significant increase in the nuclear-to-cytoplasmic ratio of NFAT1/2 in Gαq-R183Q ECs, with minimal effects in Gαq-WT cells (Fig. 5A, B). Strikingly, Western blot analysis showed clear FK506-mediated restoration of phosphorylation levels of NFAT1-S326, NFAT1-S54, and NFAT2-S294 specifically in Gαq-R183Q mutant cells, without altering their levels in Gαq-WT cells (Fig. 5C, D). Moreover, functional assays demonstrated that FK506 slightly improved wound healing capacity (Fig. 5 E, F) and partially rescued angiogenic capacity in Gαq-R183Q ECs, as evidenced by increased sprouting number and length in sprouting assays (Fig. 5G, H). These findings indicate that the calcineurin/NFAT signaling axis is important for endothelial migration and sprouting and that its dysregulation may present a potential therapeutic target.

**Figure 5.**
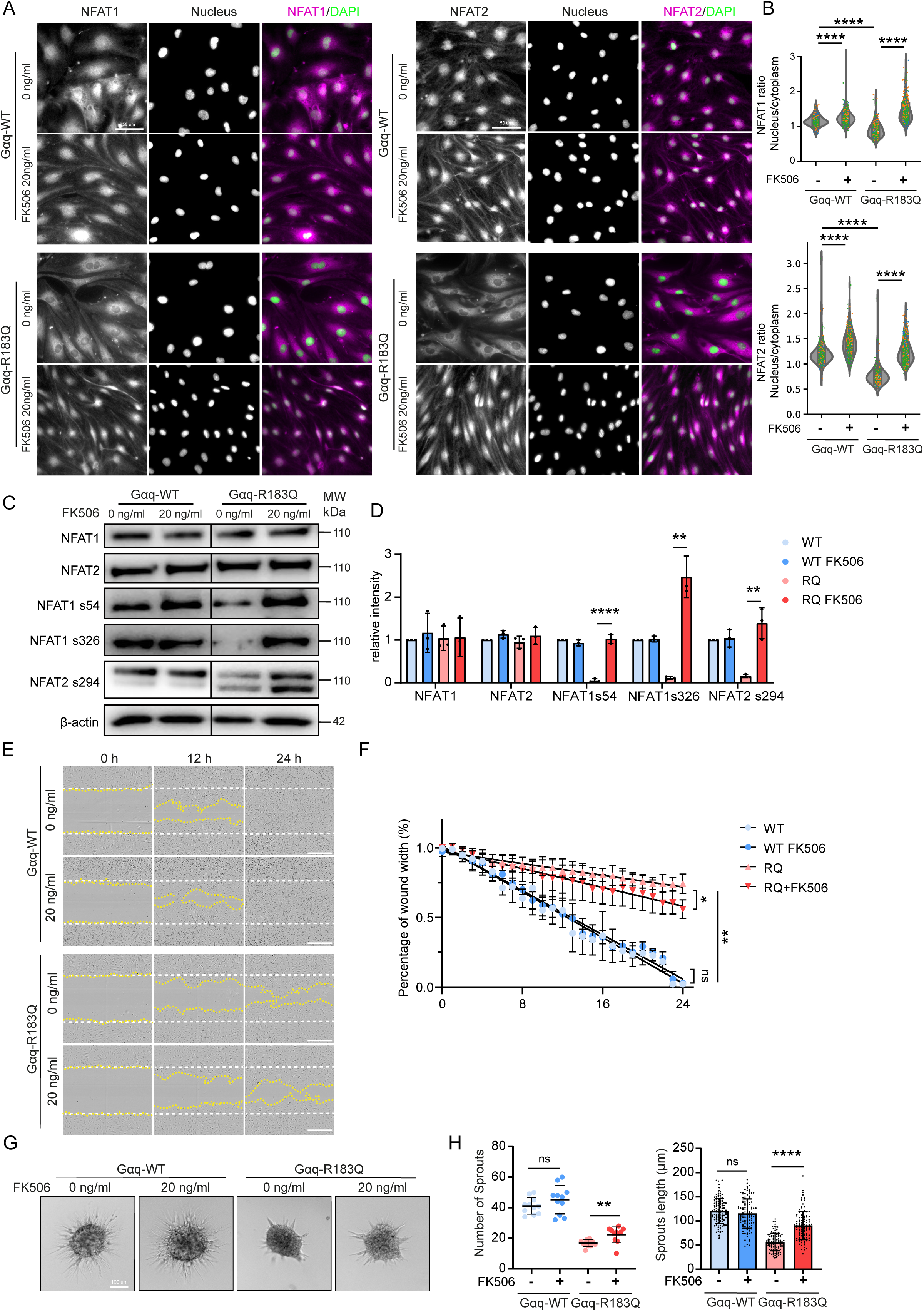
Inhibition of calcineurin activity improves functions of Gαq-R183Q ECs. (A) Immunofluorescence analysis of Gαq-WT and Gαq-R183Q cells treated with FK506 (20 ng/ml) for 24 hours, stained for NFAT1 (magenta, left), NFAT2 (magenta, right) and DAPI (green; nuclei). Scale bars: 50 μm (B) Quantification of the nuclear-to-cytoplasmic ratio of NFAT1/2 fluorescence intensity in Gαq-WT and Gαq-R183Q cells following FK506 treatment (20 ng/ml, 24 hours) using automated image processing and segmentation. Data analysed by one-way ANOVA with Tukey’s post hoc test; mean ± SD (n=5 independent experiments; N = from left to right = 249, 449, 177, 271 cells for NFAT1, N = from left to right = 251, 510, 267, 330 cells for NFAT2). (C) Western blot analysis of NFAT1, NFAT2, NFAT1-S54, NFAT1-S326, NFAT2-S294, and β-actin (top to bottom) in Gαq-WT and Gαq-R183Q cells treated with FK506 (20 ng/ml) for 24 hours. β-actin was used as a loading control. (D) Statistical analysis of immunoblot band intensities. Data analysed by one-way ANOVA with Tukey’s post hoc test; mean ± SD (n=3 independent experiments). (E) Representative phase-contrast images of Gαq-WT and Gαq-R183Q cells treated with FK506 (20 ng/ml) for 24 hours in a scratch-wound assay (t = 12 and 24 hours after scratch). The white dotted line indicates the initial wound area; the yellow dotted line marks the remaining unclosed wound area. Scale bars: 100 μm. (F) Quantification of wound width over time using Incucyte ZOOM™ software. Data are shown as mean ± SD (n=3 independent experiments). (G) Representative widefield images from spheroid-based sprouting assays of Gαq-WT and Gαq-R183Q cells treated with FK506 (20 ng/ml) for 24 hours after VEGF stimulation. Scale bar: 100 μm. (H) Dot plots showing cumulative sprout length and sprout number. Statistical analysis was performed using one-way ANOVA with Tukey’s multiple comparisons test; mean ± SD (n=3 independent experiments; N from left to right = 11, 10, 10,11 spheroids).

### Depletion of DSCR1 in Gαq-R183Q ECs restores NFAT phosphorylation, endothelial migration and angiogenic sprouting

The expression of DSCR1.4 can inhibit excessive calcineurin/NFAT signaling^35,36^, through which it controls endothelial-driven angiogenesis and inflammatory responses^38,39^. To assess the potential usefulness of DSCR1 as a molecular target for therapies in CMs, we next determined its importance for Gαq-R183Q-ECs. To achieve this, DSRC1 expression was depleted using lentiviral shRNA in Gαq-WT and Gαq-R183Q ECs, resulting in efficient silencing of DSCR1.1 and DSCR1.4 in Gαq-R183Q ECs (Fig. 6A). DSCR1 depletion significantly increased phosphorylation levels at NFAT1-S54, NFAT1-S326, and NFAT2-S294 in Gαq-R183Q ECs, but did not further increase their levels in Gαq-WT cells (Fig. 6B). DSCR1 knockdown promoted nuclear translocation of both NFAT1 and NFAT2 in Gαq-R183Q-expressing cells (Fig. 6C, D). Scratch wound assays demonstrated that DSCR1 depletion almost completely restored endothelial migration capacity in Gαq-R183Q ECs to that of normal ECs (Fig. 6E, F). Additionally, DSCR1 knockdowns strongly enhanced the angiogenic capacity of Gαq-R183Q ECs in spheroid-based sprouting assays (Fig. 6G, H). These results demonstrate that the depletion of DSCR1.4 from Gαq-R183Q endothelial cells relieves its negative feedback potential on the calcineurin/NFAT pathway, leading to a significantly improved endothelial functions. These findings indicate that if the calcineurin-NFAT-DSCR1.4 signaling axis could be specifically targeted pharmacologically it may provide a promising strategy to treat CMs.

**Figure 6.**
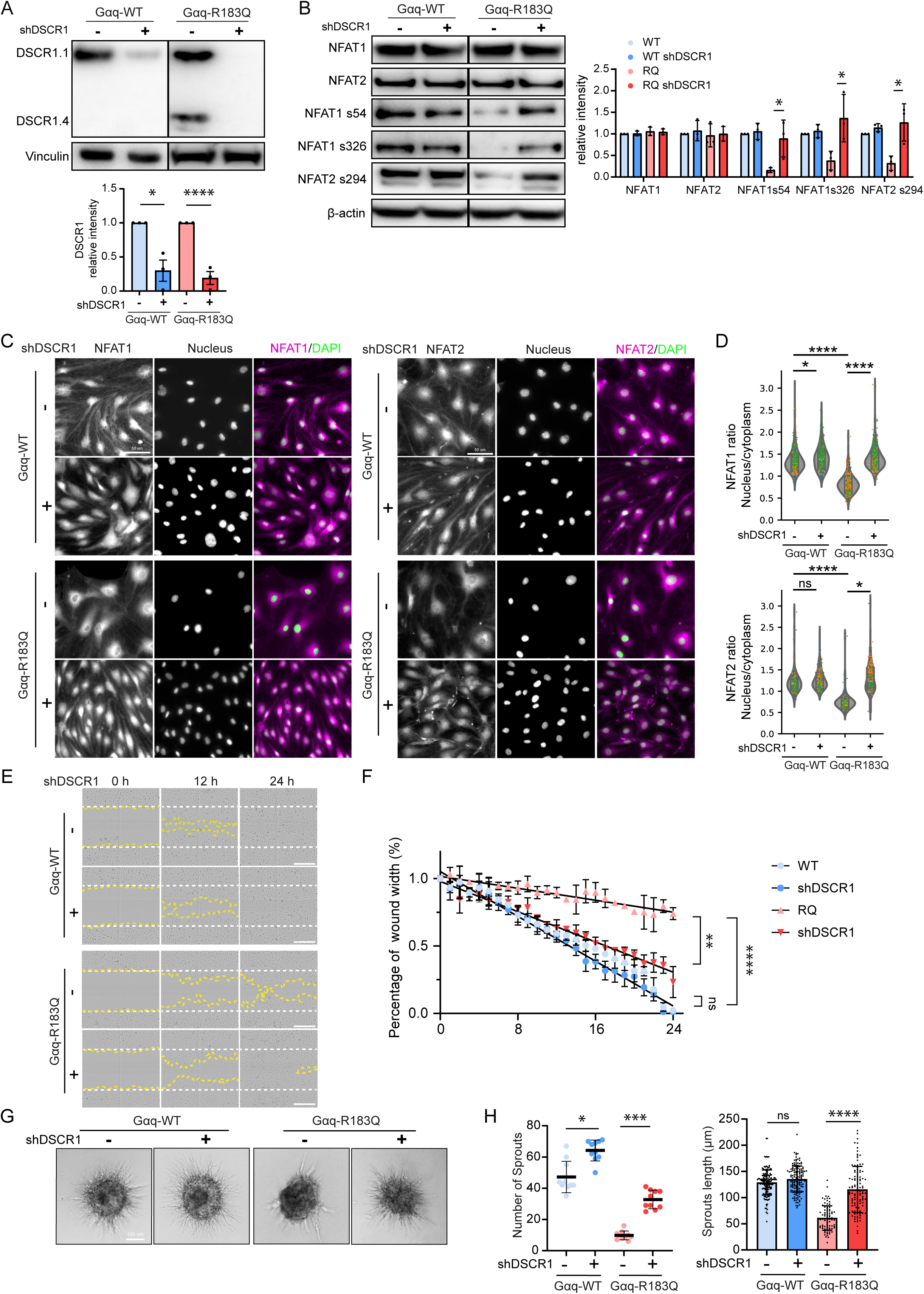
Depletion of DSCR1 in Gαq-R183Q ECs restores NFAT phosphorylation, endothelial migration and angiogenic sprouting. (A) Western blot analysis of DSCR1.1, DSCR1.4, and β-actin (top to bottom) in Gαq-WT and Gαq-R183Q cells transduced with shControl or shDSCR1. β-actin served as the loading control. Data analysed by one-way ANOVA with Tukey’s post hoc test; mean ± SD (n=3 independent experiments). (B) Western blot analysis of NFAT1, NFAT2, NFAT1-S54, NFAT1-S326, NFAT2-S294, and β-actin (top to bottom) in Gαq-WT and Gαq-R183Q cells transduced with shControl or shDSCR1. β-actin served as the loading control. Statistical analysis of immunoblot band intensities. Data analysed by one-way ANOVA with Tukey’s post hoc test; mean ± SD (n=3 independent experiments; N from left to right = 276, 510, 284, 265 cells for NFAT1, N from left to right = 178, 248, 188, 228 cells for NFAT2). (C) Immunofluorescence images of Gαq-WT and Gαq-R183Q cells transduced with shControl or shDSCR1. Cells were stained for NFAT1 (magenta, left), NFAT2 (magenta, right), and DAPI (green; nuclei). Scale bars: 50 µm. (D) Quantification of nuclear-to-cytoplasmic fluorescence intensity ratios for NFAT1 (top) and NFAT2 (bottom) in the indicated cell lines. Data analysed by one-way ANOVA with Tukey’s post hoc test; mean ± SD (n=5 independent experiments). (E) Phase-contrast time-lapse images of scratch-wound assays in Gαq-WT and Gαq-R183Q cells transduced with shControl or shDSCR1. (t=0, 12 and 24 h after scratch). The while dotted line shows the initial scratch area. The yellow dotted line highlight the unclosed wound area. Scale bars: 100 μm. (H) Wound width quantification over time using Incucyte ZOOM™ software. Data shown as mean ± SD (n=3 independent experiments). (G) Widefield microscopy images of spheroid sprouting assays in Gαq-WT and Gαq-R183Q cells transduced with shDSCR1, 16 hours post-VEGF stimulation. Scale bar: 100 μm. (H) Cumulative sprout length (right) and sprout number (left) quantified from spheroid assays. Data from n=3 independent experiments; N from left to right = 10, 10, 9, 10 spheroids; one-way ANOVA with Tukey’s post hoc test.

## Discussion

Since the identification of the p.R183Q substitution in the GNAQ gene in SWS and nonsyndromic CMs, studies focused on understanding its consequence for pathogenesis, signaling pathways, and endothelial dysfunction ^7,31,40^. In this study, we employed Gαq-knockout human dermal endothelial cells (ECs) rescued with either Gαq-R183Q or Gαq-WT to systematically map signaling in CMs through phosphoproteomic and proteomic profiling. We present, for the first time, a comprehensive comparison of the relative phosphoproteome between Gαq-WT and Gαq-R183Q endothelial cells. Our analysis identified the PI3K/PTEN and calcineurin/NFAT pathways as the most significantly dysregulated in Gαq-R183Q ECs. The Gαq-R183Q mutation activates multiple downstream pathways, including the canonical PLCβ pathway towards IP_3_ and DAG^41^. IP_3_ stimulates calcium release and the calcineurin/NFAT pathway^31^. Our current study demonstrated downregulation of phosphosites relevant to activation of the calcineurin/NFAT pathway in Gαq-R183Q ECs (NFAT1-S326, NFAT1-S54, and NFAT2-S294) (Table S1).

While the NFAT target DSCR1 has previously been implicated in angiogenesis, tumor growth, and megakaryopoiesis via calcineurin/NFAT pathway^42–44^, its role in Gαq-R183Q-driven CMs via calcineurin/NFAT regulation has not yet been studied. Previous transcriptomic studies indicated that the PLCβ/PIP_2_/IP_3_ pathway stimulates the calcineurin/NFAT pathway, leading to enhanced DSCR1.4 expression in Gαq-R183Q mutated CMs^31^. Here, we confirm that the calcineurin-NFAT pathway is not only hyperactive in Gαq-R183Q ECs, but also critically regulates endothelial migration and sprouting motility through upregulating expression of DSCR1.4. The elevated DSCR1.4 levels in Gαq-R183Q ECs correlated with impaired nuclear import of NFAT1/2. Since DSCR1.4 functions as an endogenous calcineurin inhibitor, it points to deregulated self-limiting signaling in CMs^45^ ^46^. Our findings thus demonstrate sustained activation of a negative feedback loop in Gαq-R183Q ECs which affects endothelial angiogenic activities. Interestingly, repression of NFAT signaling has been reported as a driver of highly angiogenic endothelial states in infantile hemangiomas^47^, a childhood vascular tumor that is driven by a hyperactivating of p.Q209L mutation in *G*NAQ^48^.

The PI3K/PTEN pathway is a well-established regulator of endothelial behavior in vascular anomalies, including CMs^49–51^. Our data confirm that the PI3K/PTEN pathway represents one of the regulatory pathways in Gαq-R183Q mutated CMs. Previous studies have reported phosphorylation changes in downstream targets of the PI3K/PTEN pathway, such as AKT, TSC1/2, mTOR, and EIF4EBP1 in CMs and other vascular malformations which is consistent with our analysis (Figure 2D, Table S1). Another key player implicated in CMs is Angiopoietin-2 (ANGPT2), which is transcriptionally upregualted in Gαq-R183Q-overexpressing endothelial cells^20,52^ and has been proposed as a biomarker for CMs. The current phosphoproteomic analysis revealed no differential ANGPT2 phosphorylation levels in Gαq-R183Q ECs. This is likely explained by the notion that phosphoproteomics primarily detects intracellular signaling events, whereas Angiopoietin-2 functions as a secreted protein ^53^.

Several FDA-approved calcineurin inhibitors, including cyclosporin A, voclosporin, pimecrolimus, tacrolimus (FK506), and basiliximab are widely used in transplantation, autoimmune disorders, and inflammatory diseases^54,55^. In this study, we focused on FK506 and its effects on Gαq-R183Q-mutated ECs. Our findings demonstrate that FK506 suppresses calcineurin/NFAT signaling by restoring NFAT phosphorylation and enhancing angiogenic capacity in Gαq-R183Q CMs. However, while FK506’s calcineurin inhibition shows therapeutic potential for mitigating NFAT-driven vascular dysfunction, its efficacy in rescuing endothelial migration and sprouting growth was less pronounced compared to the genetic depletion of DSCR1.4. This difference may result from reduced expression of FK506-binding proteins (FKBPs) in Gαq-R183Q endothelial cells. Phosphoproteomic analysis showed an approximately 1-fold decrease in FKBPs levels (including FKBP1A, FKBP11, and FKBP3) in mutant cells compared to controls. FKBPs play critical roles in endothelial signaling, angiogenesis, and vascular integrity by modulating NF-κB and mTOR/AKT pathways^56,57^. Since FK506 requires FKBPs to inhibit the calcineurin/NFAT axis, diminished FKBP expression likely attenuates the drugs inhibitory efficacy, resulting in weaker functional effects compared to DSCR1.4 knockdown. Moreover, calcineurin inhibitors may only partially suppress the dysregulated signaling in endothelial GNAQ R183Q-induced capillary malformations, as parallel signal transduction pathways can remain active and continue to drive lesion development.

In summary, our study utilized a Gαq-R183Q-mutated CMs *in vitro* model to delineate the cellular and functional consequences of this mutation in endothelial cells. We identified the calcineurin-NFAT-DSCR1.4 signaling axis as a critical regulator of endothelial functions in Gαq-R183Q CMs. This signal transduction pathway is amenable for modulation through both DSCR1.4 silencing and calcineurin inhibitors, highlighting it as a potential therapeutic target for the development of selective therapies in CMs and SWS patients. Repurposing existing drugs targeting the calcineurin pathway represents a potentially effective and cost-efficient strategy for developing new treatment options for CMs. Moreover, the presence of DSCR1.4 in capillary endothelium may serve as a potential biomarker for CMs.

## Methods and Materials

### Antibodies and reagents

The following antibodies and dyes were used for immunostaining (IF) and immunoblotting (WB) in this study. For immunofluorescence, we used Alexa Fluor 647-conjugated mouse monoclonal anti-human VE-cadherin (CD144, clone 55-7H1, BD Biosciences, Cat# 561567; 1:200), and PromoFluor-488 Phalloidin for F-actin staining (Promokine, Cat# PK-PF488P-7-01; 1:200). Alexa Fluor 594 chicken anti-rabbit (Thermo Fisher Scientific, Cat# A21442; 1:500) and Alexa Fluor 647 chicken anti-mouse (Thermo Fisher Scientific, Cat# A21463; 1:500) were used as secondary antibodies. Nuclei were stained with DAPI (Thermo Fisher Scientific, Cat# D3571, 1:1000). For immunoblotting, we used rabbit monoclonal anti-GNAQ (clone EPR17149, Abcam, Cat# ab199533; 1:1000), rabbit polyclonal anti-alpha-tubulin (Proteintech, Cat# 11224-1-AP; 1:10,000), mouse monoclonal anti-GFP (clone B-2, Santa Cruz Biotechnology, Cat# sc-9996; 1:1000), rabbit polyclonal anti-NFAT1 s54 (Thermo Fisher Scientific, Cat# 44-944G; 1:1000), rabbit polyclonal anti-NFAT1 s326 (abcepta, Cat# AP61143; 1:1000), rabbit polyclonal anti-NFAT2 s294 (Amerigo Scientific, Cat# STJ9118; 1:1000). Additional antibodies included anti-DSCR1 (Sigma-Aldrich, Cat# D6694; 1:1000 for WB), anti-NFAT1 (Cell Signaling Technology, Cat# 5861-T; 1:1000 for WB, 1:200 for IF), and anti-NFAT2 (Abcam, Cat# ab25916; 1:1000 for WB, 1:200 for IF). For WB secondary antibodies conjugated to horseradish peroxidase (HRP) were obtained from BioRad.

### Cell culture and plasmids

CRISPR/Cas9-mediated *GNAQ* knockouts of immortalized human dermal microvascular endothelial cell lines (HDMEC; ATCC CRL4025) were generated as follows. Briefly, annealed gRNA-FW and gRNA-REV oligonucleotides (Supplementary Table 4) were inserted into the pSpCas9(BB)-2A-GFP plasmid (Addgene #48138) and transfected into wild-type HDMECs. GFP-positive cells were isolated by fluorescence-activated cell sorting (FACS), expanded from single colonies, and cryopreserved until further use. Plasmid encoding intramolecularly mTurquoise-tagged Gαq, a validated fully functional fluorescent reporter^58^, was a kind gift of Dr. Joachim Goedhart. Gαq-R183Q-mTurquoise was generated using site-directed mutagenesis. Gαq-WT-mTurquoise and Gαq-R183Q-mTurquoise inserts were cloned into a self-inactivating lentiviral pLV-CMV-ires-puro vector. Lentiviral particles were produced in HEK293T cells, which were transiently transfected with third-generation packaging constructs and the lentiviral expression vector of interest using Trans-IT LTI (Mirus). Subsequently, *GNAQ* KO HDMEC were transduced with lentivirus produced from either pLV-Gαq-WT-mTurquoise or pLV-Gαq-R183Q-mTurquoise plasmids, and selected with puromycin (sigma, 1 µg/mL) to establish stable cell lines constitutively expressing either wild-type Gαq or mutant Gαq-R183Q. shControl (shC002) lentiviral constructs and the shRNA in the lentiviral pLKO.1 backbone targeting DSCR1 (TRCN0000019844) were from the Sigma-Aldrich mission library. All cell lines were cultured on gelatin-coated dishes in Endothelial Cell Growth Medium (Promocell, Cat# C-22111) with 1% Penicillin/Streptomycin (Penicillin: 100U/mL, streptomycin: 100µg/mL) at 37°C in a humidified atmosphere with 5% CO₂.

### Western blot analysis

Cells were lysed in 1X Laemmli buffer (4% SDS, 48% Tris 0.5 M pH6.8, 20% glycerol, 18% H2O and bromophenol blue) containing 4% β-mercaptoethanol. Lysates were boiled at 95°C for 5 minutes to denature proteins. Proteins were separated on 10% SDS-PAGE gels using running buffer (25 mM Tris-HCl, pH 8.3, 192 mM glycine, and 0.1% SDS), and subsequently transferred to ethanol-activated PVDF membranes via wet transfer in blotting buffer [25 mM Tris-HCl, pH 8.3, 192 mM glycine, and 20% (v/v) ethanol]. Membranes were blocked with 5% non-fat milk in Tris-buffered saline (TBS) for 1 hour at room temperature. Blots were incubated overnight at 4°C with primary antibodies diluted in 5% milk in TBS containing 0.1% Tween-20 (TBST). After washing, membranes were incubated with HRP-conjugated secondary antibodies for 45 minutes at room temperature. Following additional washes with TBS, protein bands were detected using enhanced chemiluminescence (ECL) reagents (SuperSignal West Pico PLUS, Thermo Fisher, Cat #34580) and visualized with an ImageQuant LAS 4000 (GE Healthcare) imaging system. Band intensities were quantified using the Gel Analyzer plugin in ImageJ.

### Immunofluorescence stainings and microscopy

For immunofluorescence staining, cells grown on fibronectin-coated (1:400) coverslips were fixed in 4% paraformaldehyde (PFA) diluted in PBS supplemented with 1 mM CaCl₂ and 0.5 mM MgCl₂ (PBS++) for 10 minutes at room temperature. After fixation, cells were washed twice with PBS++, permeabilized with 0.5% Triton X-100 in PBS for 5 minutes at room temperature, and washed again twice with PBS. Blocking was performed in 2% BSA in PBS. Cells were then incubated with primary antibodies against VE-cadherin and F-actin for 1 hour in the dark. Following incubation, coverslips were washed with PBS and mounted onto microscope slides using Mowiol 4-88 (Calbiochem, #475904) containing DABCO (Sigma-Aldrich, D27802) as an antifade agent. Images were acquired using a Nikon Eclipse TI widefield microscope equipped with a Lumencor SOLA SE II light source, standard CFP, GFP, mCherry, or Cy5 filter cubes, a 60x Apo TIRF oil objective (NA 1.49), and an Andor Zyla 4.2 plus sCMOS camera.

To determine nucleus-to-cytoplasm ratios a custom Python script was used (Github Repository: https://github.com/Jintram/Analysis_nucleus_cyto_ratio_static). The script segments the nuclei based on the DAPI channel (using an Otsu threshold after median filtering) and determines a narrow ring around the nucleus to characterize the cytoplasm (using image dilation). For each cell, the NFAT nucleus-to-cytoplasm ratio is calculated by dividing the mean background corrected (subtraction of mode) NFAT intensity in the nuclear region by that in the cytoplasmic ring. After automated analysis, the output was reviewed manually to remove mis-segmented cells and artifacts.

### Immunofluorescence of patient samples

Formalin-fixed, paraffin-embedded (FFPE) CM biopsy blocks were obtained from the Department of Dermatology at a tertiary Vascular Anomalies Center, Amsterdam University Medical Centers (Amsterdam UMC), The Netherlands. The study adhered to the Declaration of Helsinki, and written informed consent was obtained from all patients. The study was approved by the Medical Ethics Committee from the Amsterdam UMC (case number NL75128.018.20), was registered at the National Trial Register in the Netherlands on February 23, 2021 (trial identification NL9295) and its direct outcome have previously been published^9^. Sections of 4 μm thickness were mounted on microscope slides, dewaxed using xylene (#28979.294, VWR Chemicals), and rehydrated through graded alcohols to water. Antigen retrieval was performed in Tris-EDTA buffer (pH 9.0) using a microwave-based protocol. Slides were blocked in 2% BSA in PBS-Tween (#P1379, Sigma Aldrich) for 1 hour. Immunofluorescence staining was conducted using primary antibodies against NFAT1 (1:100), NFAT2 (1:100), DSCR1 (1:100) and VE-cadherin (1:100). Slides were incubated with primary antibodies overnight at 4 °C, followed by two washes with PBS-Tween and incubation with appropriate secondary antibodies for 1 hour at room temperature. After additional PBS-Tween washes, slides were mounted with Mowiol 4-88 containing DABCO as an antifade agent. Imaging was performed using a Leica STELLARIS 8 confocal microscope (Leica Microsystems, Wetzlar, Germany) equipped with a 63x oil immersion objective.

### Scratch assays

For the scratch assays in Fig. 1, cells were seeded onto 12-well plates coated with 5 µg/ml fibronectin. Once a confluent monolayers were established, two perpendicular scratches per well were made using a sterile 200 µl pipette tip. Detached cells were removed by washing with PBS++, after which the cells were cultured in EGM-2 medium. Plates were mounted on an inverted Nikon Eclipse TI microscope equipped with an Okolab cage incubator and a humidified CO₂ chamber maintained at 37°C and 5% CO₂. Live-cell imaging was performed for 17 hours at 10-minute intervals using phase-contrast microscopy with a 10× CFI Achromat DL dry objective (NA 0.25) and an Andor Zyla 4.2 plus sCMOS camera. Images were processed for display using an unsharp mask filter and adjusted for brightness and contrast in ImageJ. The wound area was quantified by measuring the scratch surface using the freehand tool in ImageJ. For experiments in Fig. 5 and 6, cells were cultured in 96-well Incucyte ImageLock plates (Sartorius, #4806) and analyzed using the Incucyte live-cell imaging system. For the migration assay, cells were seeded into the plates and grown to approximately 90% confluency. A uniform scratch was created in each well using the Incucyte WoundMaker tool. Images were captured every 30 minutes over a 24-hour period. Image analysis was performed using the Incucyte Base Analysis Software in combination with the Incucyte Plate Map Editor.

### Sprouting angiogenesis assays

For sprouting angiogenesis assays, cells were resuspended in EGM-2 medium containing 0.1% methylcellulose (4000 cP, Sigma). To generate spheroids, 750 cells in 100 µl of methylcellulose medium were seeded into each well of a U-bottom 96-well plate and incubated overnight to allow spheroid formation. The resulting spheroids were collected, resuspended in a 1.7 mg/ml collagen type I rat tail mixture (IBIDI), and plated into glass-bottom 96-well plates, which were then incubated at 37°C to allow gel polymerization. After the collagen gel had solidified, spheroids were stimulated with 50 ng/ml VEGF (Thermo Fisher Scientific) to induce sprouting overnight, as previously described^59^. Images were acquired using the EVOS M7000 imaging system with a 10× objective. Sprout number and length were quantified using the NeuronJ plugin in ImageJ.

### SILAC preparation and LC-MS/MS measurement

Gαq-KO HDMECs expressing either Gαq-WT-mTurquoise2 or Gαq-R183Q-mTurquoise2 were passaged five times in SILAC EBM-2 culture medium (Lonza), lacking arginine and lysine, and supplemented with EGM-2 BulletKit and 2% FCS. Cells were cultured in SILAC medium using either ‘heavy’ medium with isotopically labeled Arg-10 and Lys-8 or ‘light’ medium with unlabeled Arg-0 and Lys-0. Lys0 (L-Lysine•2HCl ^12^C_6_^14^N_2_, REF = 88429, Cat #WB322018, Thermofisher), Arg0 (L-Arginine•HCl ^12^C_6_^14^N_4_, REF = 88427, Cat # WA321431, Thermofisher), Lys8 (L-Lysine•HCl ^13^C_6_^15^N_2_ REF= 211603902, Cat # 211CXN-LysI-708-01, Silantes), Arg10 (L-Arginine•HCl ^13^C_6_^15^N_4,_ REF = 201603902, Cat # 201CXN-ArgI-2009-01, Silantes) were dissolved according to manufacturer instructions.

For each condition, cells were plated in two 15 cm dishes and cultured to high confluency. Cells were harvested by scraping into 1.5 mL lysis buffer containing 8 M urea (MP Biomedicals, Cat# 821527), 1 M ammonium bicarbonate (Fluka, Cat# 09830), 10 mM Tris(2-carboxyethyl)phosphine hydrochloride (TCEP) (Sigma, Cat# 75259), 40 mM 2-chloroacetamide (CAA; Sigma, Cat# C0267), 1% (v/v) phosphatase inhibitor cocktail 2 (Sigma, Cat# P5726), 1% (v/v) phosphatase inhibitor cocktail 3 (Sigma, Cat# P0044), and one tablet of EDTA-free protease inhibitor cocktail (Roche, Cat# 12273700). Lysates were immediately combined from the two dishes per condition to create label pairs, with a label swap included as a control for label dependency. Samples were sonicated to shear DNA and homogenize the lysate, then diluted fourfold with 1 M ammonium bicarbonate and digested overnight with trypsin (Worthington Biochemical Corp, Cat# LS003750) at a 50:1 protein-to-trypsin ratio, in the presence of 1 mM CaCl₂ at 37°C on a shaker at 1000 rpm.

Digested peptides were cleaned using C18 columns (Waters, 3 CC, C18 200 mg cartridge) and eluted with 1 mL of 80% acetonitrile (ACN). For phosphopeptide enrichment, samples were acidified with trifluoroacetic acid (TFA) to a final concentration of 6%, then incubated with calcium titanate (CaTiO₃) powder (Alfa Aesar, 325 mesh) at a ratio of 10:1 (protein:powder) for 10 minutes at 40°C on a shaker. Following phosphopeptide binding, the CaTiO₃ powder was washed to remove non-phosphorylated peptides. The supernatant was removed, and 1 mL of wash solution (80% ACN, 1% TFA) was added. Samples were vortexed, centrifuged (500 rpm, 30 seconds), and the supernatant discarded. This wash cycle was repeated three times. For elution, 300 µL of 5% ammonia solution was added to the beads, and the samples were incubated for 10 minutes at room temperature (RT) on a heater-shaker at 800 rpm. After centrifugation, the phosphopeptide-containing supernatant was transferred to a new 1.5 mL tube and acidified with 20 µL of 20% formic acid. Samples were further cleaned using C18 StageTips and fractionated using high-pH reverse-phase fractionation into three fractions. The samples were vacuum-dried and then resuspended in 10 µL of 0.1% formic acid. All samples were vacuum-dried. Samples were resuspended in 10 µL of 0.1% formic acid, and the first two fractions were analyzed on an Orbitrap Eclipse (Thermo Fisher Scientific) using a 240-minute gradient on a self-packed 50 µm C18 column. The mass spectrometry method employed a data-dependent MS2 approach with 240,000 resolution (400–1600 m/z range), HCD fragmentation at 30% NCE, 100% AGC, dynamic injection time, a cycle time of 1 second, and a 60-second dynamic exclusion. Two FAIMS settings at -45 V and -65 V were used during acquisition.

### SILAC Proteomics Data Analysis

Raw mass spectrometry data were analyzed using MaxQuant (version 1.6.3.4) against the complete *Homo sapiens* proteome (taxonomy ID: 9606) obtained from UniProtKB. Default MaxQuant parameters were applied, with specific settings for SILAC labeling using Arg10 and Lys8. Phosphorylation of serine, threonine, and tyrosine residues (Phospho (STY)) was included as a variable modification, in addition to methionine oxidation. Trypsin was specified as the protease, allowing up to two missed cleavages. MaxQuant output files, “proteinGroups.txt” and “Phospho (STY)Sites.txt,” were used for downstream analysis in R (version 4.0.4). Proteins and phosphopeptides were initially filtered to remove contaminants, reverse sequences, and proteins identified by fewer than two unique peptides. For phosphopeptides, additional filtering was performed based on the localization score. Duplicate protein entries were resolved by merging data according to gene names and protein IDs. For each identification and sample pair, the reported normalized SILAC ratios were extracted. Ratios were log2-transformed and centered to facilitate comparative analysis (Supplementary Figure 3). Significance thresholds were set at ±2 standard deviations from the mean, highlighting the top 5% of significant changes. Missing values were imputed using the minimum observed ratio, adjusted by a small constant, to enable their visualization in downstream analyses. The mass spectrometry proteomics data have been deposited to the ProteomeXchange Consortium via the PRIDE^60^ partner repository with the dataset identifier PXD064187.

### Ingenuity Pathway Analysis

All phosphopeptides and their corresponding fold-change values were uploaded into Ingenuity Pathway Analysis (IPA, Qiagen, free trial version) to identify enriched upregulated and downregulated pathways, as well as potential upstream regulators. Z-scores and p-values were calculated in IPA using Fisher’s exact test^61^. For visualization, the top 15 significant pathways were plotted for Gαq, while all significant pathways were plotted for Gαq-R183Q. IPA graphs were generated in R.

### Statistical analysis

All data and statistical analyses were performed using GraphPad Prism version 6. Bar graphs represent mean ± standard deviation (SD), while violin plots indicate the median and quartiles. Data were assessed for normality using the D’Agostino-Pearson normality test. For data with a normal distribution, a two-tailed paired Student’s t-test was used to compare two groups. For comparisons involving more than two groups, one-way or two-way analysis of variance (ANOVA) was performed, followed by post hoc tests as specified in the figure legends. The specific statistical tests used are indicated in the corresponding figure descriptions. P-values are represented by asterisks and defined as follows: ns, not significant; *P < 0.05; **P < 0.01; ***P < 0.001; ****P < 0.0001.

## Acknowledgements

This publication is part of the RepairQ project with file number era4healthcvd-122 of the ERA for Health Cardinnov research program which is financed by the Dutch Research Council (NWO) and the Dutch Heart Foundation. This research was further financially supported by the Netherlands Organization of Scientific Research (NWO OCENW.KLEIN.281) and the AMC Foundation.

## Data availability

Mass spectrometry proteomics data are available via ProteomeXchange with identifier PXD064187.

## Figure legends

**Supplementary Figure 1. Immunofluorescence staining of skin biopsies from CM patients.** (A) Representative images showing DSCR1 staining (magenta) in a skin biopsy. Endothelial cells in blood vessels were identified by VE-cadherin staining. Nuclei were stained with DAPI. Scale bar = 100 µm. (B) Representative images showing NFAT1-S326 (left) and NFAT2-S294 (right) staining in skin biopsies. Endothelial cells were marked with VE-cadherin (white), and nuclei were stained with DAPI (green). Scale bar = 50 µm.

**Supplementary Figure 2. Working model** The Gαq-R183Q mutation leads to the activation of PLCβ signalling. Activated PLCβ produces the second messengers IP₃ and DAG, which in turn activate calcineurin and protein kinase C (PKC), respectively. Once the calcineurin pathway is activated, it dephosphorylates NFAT1 and NFAT2, enabling their translocation to the nucleus where they regulate gene expression. FK506 (Tacrolimus) forms a complex with FK506 binding proteins (FKBPs), which inhibits calcineurin activity and thereby prevents NFAT1/2 dephosphorylation and nuclear translocation. Similarly, DSCR1.4 inhibits the calcineurin-NFAT pathway by blocking the dephosphorylation of NFAT1/2, thus preventing their nuclear translocation. Schematic created with Biorender.com.

**Supplementary Figure 3.** Scatterplot showing the relative phosphopeptide abundance in Gαq-WT and Gαq-R183Q ECs. The x-axis represents normalized log₂ ratios from the forward experiment, while the y-axis represents normalized log₂ ratios from the reverse experiment. Statistically significant peptides (p < 0.05) are highlighted in blue.

**Supplementary Table 1.**List of differentially detected phosphorylated sites identified in the SILAC analysis of Gαq-WT and Gαq-R183Q.

**Supplementary Table 2**. List of differentially expressed proteins identified in the SILAC analysis of Gαq-WT and Gαq-R183Q.

**Supplementary Table 3**. PTEN activation associated phosphorylated genes.

**Supplementary Table 4.** List of guide RNA used for pSpCas9(BB)-2A-GFP plasmid

## Notes

### Competing Interest Statement

The authors have declared no competing interest.

