## Supplementary figures and images for "Phosphoproteomics Maps Calcineurin-NFAT-DSCR1.4 Signaling as Druggable Axis in Gαq-R183Q–Driven Capillary Malformations"

### Supplemental Figures 1-3

Supplementary Figure 1

A

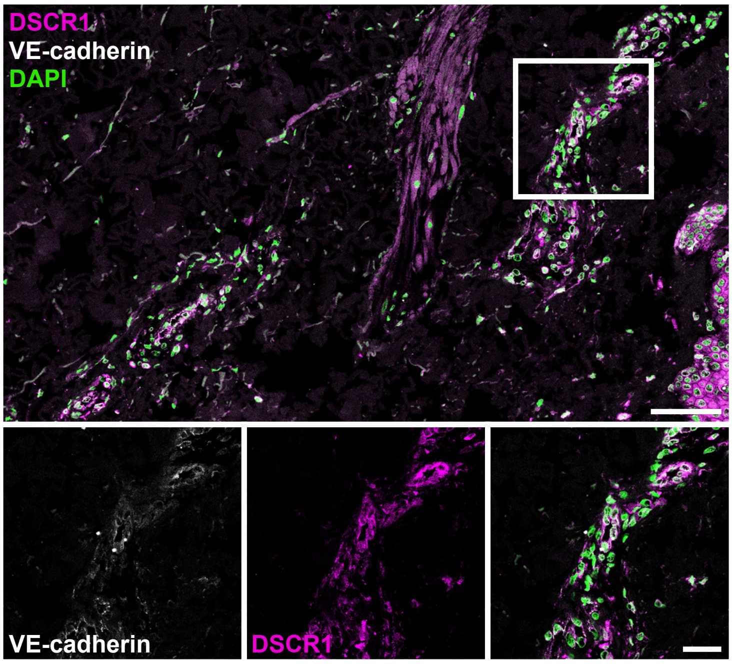


B

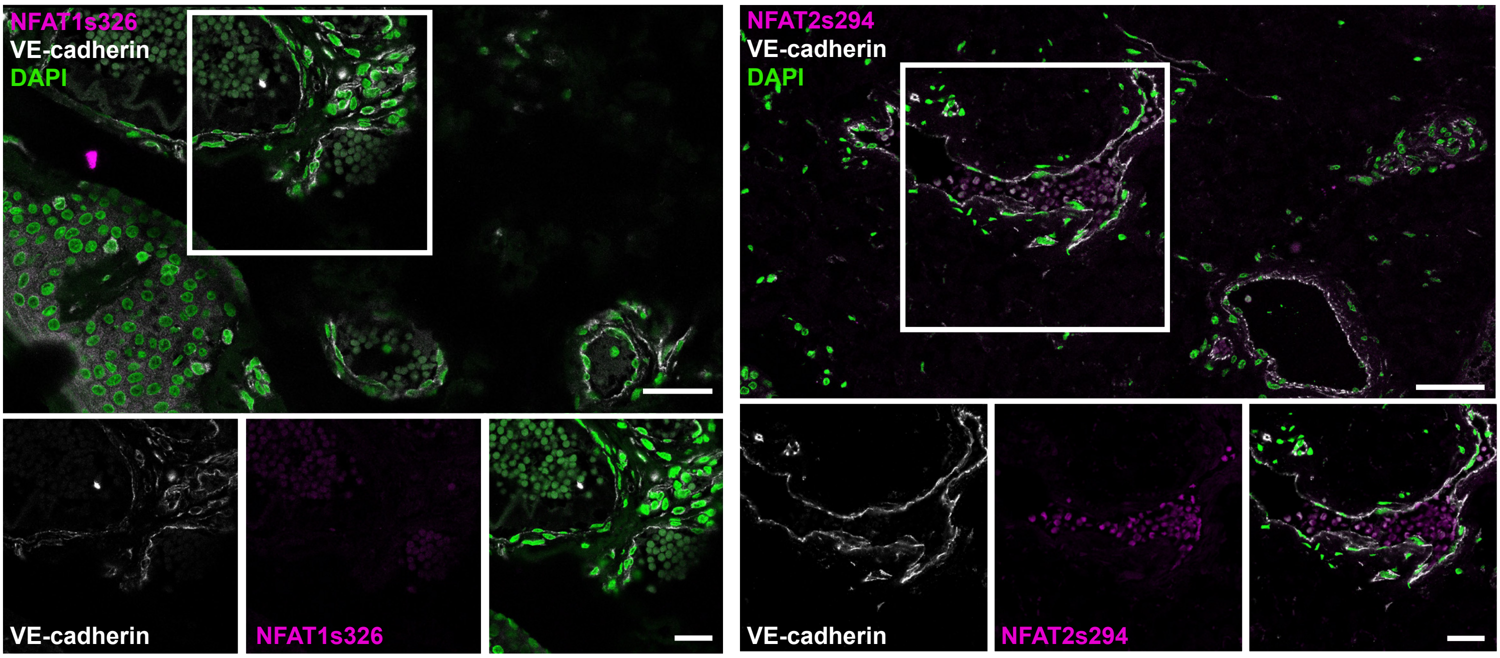

Supplementary Figure 2

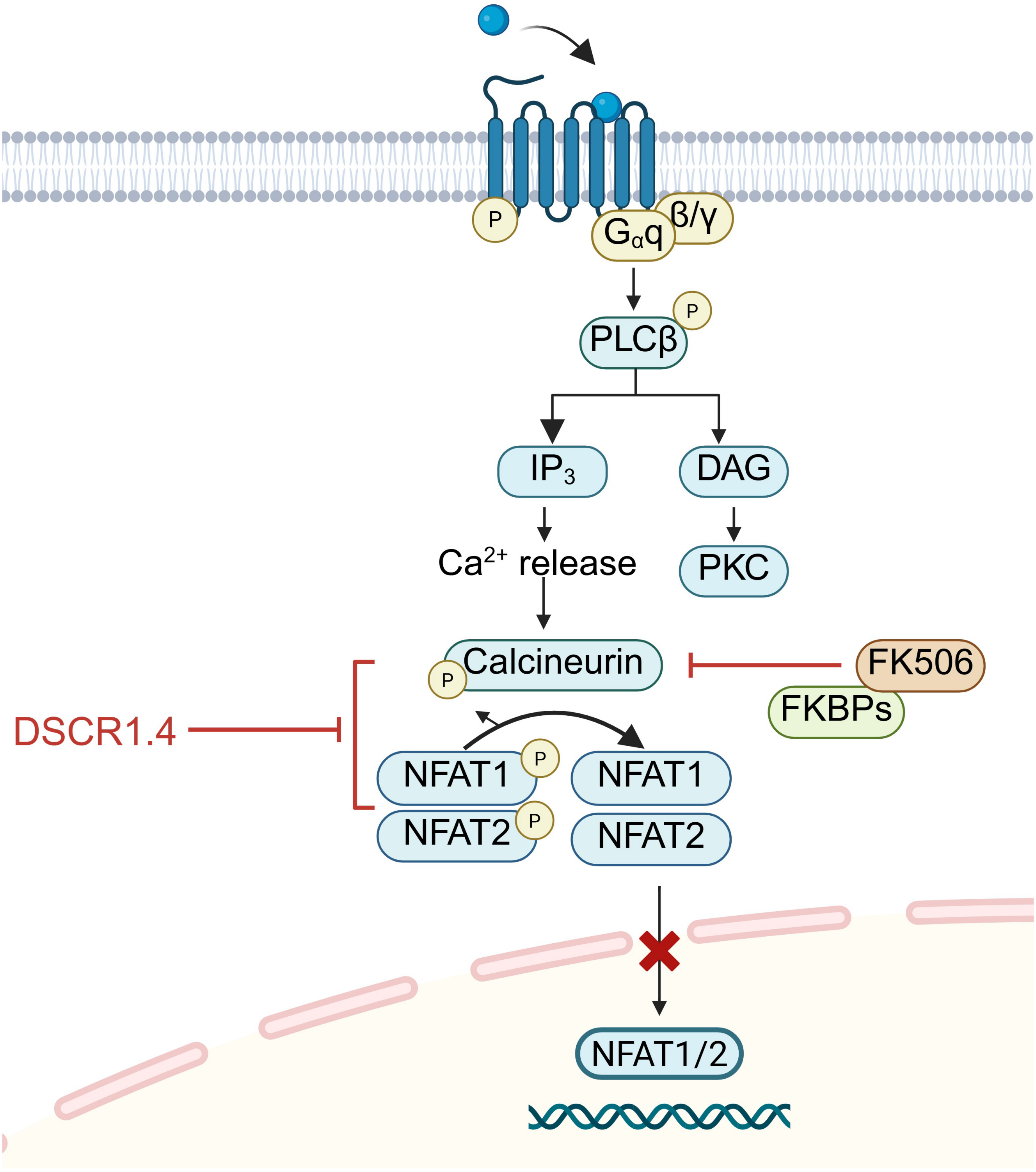

Supplementary Figure 3

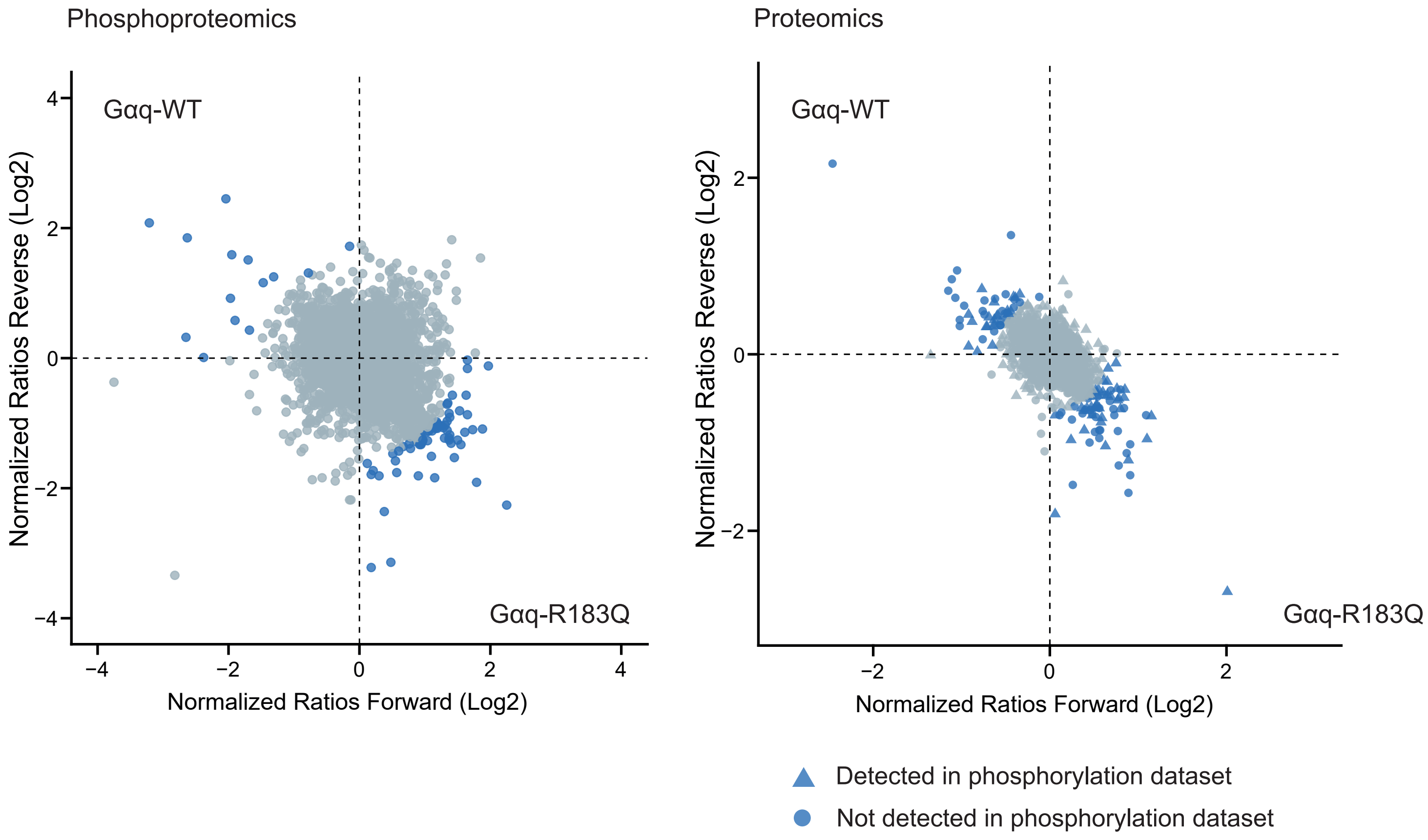
